# Discordant gene responses to radiation in humans and mice: hematopoietically humanized mice may save the day for radiation biomarker identification

**DOI:** 10.1101/558882

**Authors:** Shanaz A. Ghandhi, Lubomir Smilenov, Monica Pujol-Canadell, Sally A Amundson

## Abstract

The mouse (*Mus musculus*) is an extensively used model of human disease and responses to stresses such as ionizing radiation. As part of our work developing gene expression biomarkers of radiation exposure, dose, and injury, we have found many genes are either up-regulated (e.g. *CDKN1A, MDM2, BBC3*, and *CCNG1*) or down-regulated (e.g. *TCF4* and *MYC*) in both species after irradiation. However, we have also found genes that are consistently up-regulated in humans and down-regulated in mice (e.g. *DDB2, PCNA, GADD45A, SESN1, RRM2B, KCNN4, IFI30*, and *PTPRO*). Here we test a hematopoietically humanized mouse as a potential *in vivo* model for biodosimetry studies, measuring the response of these 14 genes one day after radiation exposure, and comparing it with that of human blood irradiated *ex vivo*, and blood from whole body irradiated mice. We found that human blood cells in the hematopoietically humanized mouse *in vivo* environment recapitulated the gene expression pattern expected from human cells, not the pattern seen from *in vivo* irradiated normal mice. The results of this study support the use of hematopoietically humanized mice as an *in vivo* model for radiation gene expression studies relevant to humans.

---

Most *in vivo* studies in which effects of external stimuli on human health are assessed require a model organism, and the mouse (*Mus musculus*) is the best studied small animal model system. A number of studies have used *Mus musculus* to identify biomarkers for a variety of diseases and biological responses, with the potential to translate them to the context of human health. In the field of radiation biology, there is a need to identify and validate markers of radiation and survival that can be used to assist emergency medical triage decisions on the basis of rapid and efficient molecular and genetic assays^1-3^. The limited availability of human samples to study biological and stress responses across a broad range of doses and times that realistically replicate real life radiological events has led to the use of *ex vivo* human and *in vivo* animal models for radiation response. Although the *ex vivo* model has been very useful to determine gene expression responses to radiation in human blood^2-4^, it is limited by culture times and conditions and lacks an *in vivo* microenvironment and tissue context. To overcome such limitations in other fields, humanized animals have been proposed to bridge the gap between *in vitro/ex vivo* models to the more realistic *in vivo* systems. One of these is the humanized NSG model^5^, in which mature human T and B cells can be developed after engraftment with human blood stem cells, making it a potentially attractive model for studying both immune development and response to stress *in vivo*.

Many researchers have addressed the need for more in depth investigations into the differences between human and rodent immunology^6^. In the last decade, there have been many studies that address and question the applicability of the mouse model to humans. One of the most notable was an extensive study on gene expression response by Seok et al in 2013^7^, in which the authors compared the response of human and mouse immune systems to wound, trauma and endotoxin exposure under comparable conditions. The best correlations between the different stress stimuli were intra-species, and the authors performed a meta-analysis on published radiation response genes in mice and humans and showed that this was also true for radiation response. From the cross-species analyses the proportion of genes that responded in the same direction in both mouse and humans was <60% across a range of doses and times^7,8^. A responding study by Takao et al^9^ in 2015 argued that the same datasets, if analyzed differently, would show that the mouse response recapitulated most of the human response, both at the gene and pathway levels. This discussion has continued on with arguments and evidence on both sides ^10-14^. Further, transcriptome-based studies comparing immune development processes such as erythropoiesis have shown that there is considerable divergence between these two species^15,16^.

In the field of radiation biology, and specifically in the area of identification of biomarkers of radiation exposure and survival based on gene expression changes in blood, we and others have identified gene panels that have the potential to be used to determine exposure to radiation in humans and model animals^2-4,8,17-28^. These studies have been effective in identifying the similarities in response between humans, mice and in some cases large animal models such as non-human primates. However, biomarker discovery and validation using human blood irradiated *ex vivo*, human blood from patients after total body irradiation, and the use of mice are limited by either lack of *in vivo* microenvironment, backgrounds of patients and their heterogeneity and species differences, respectively. Therefore, we chose to test the gene response to radiation of human blood cells that mature *in vivo* in a humanized mouse model, which provides a healthy *in vivo* microenvironment.

Humanized mouse models are modified to either express human genes or are engrafted with human cells/tissues with the goal of generating *in vivo* experimental models for pre-clinical testing and investigation of other human biological processes. In the current study we specifically chose the NSG (NOD/LtSz-scid Il2rg^−/−^) mouse model that has the advantages of a long-life span, absence of NK cells, ability to develop mature T and B cells and a functional human immune system, and radiation sensitivity^5,29,30^. We exposed the mice to total body irradiation and studied differential gene expression 24 hours later. We focused on a group of genes selected from our previous studies, some of which were expected to show a similar directional response between species and some an opposite response. We validated these response patterns using quantitative real-time PCR in irradiated non-engrafted mice and *ex vivo* irradiated human blood, in addition to the measurements made in humanized mice. Meta-analysis of previous microarray comparisons also suggested differences in the radiation responsive behavior of signaling modules centering on P53 protein family members, MYC and STAT transcription factors. These signaling and gene expression differences between humans and mice suggest that assumptions of similarity between these two mammals may not always be true and need to be individually assessed within the context of any study. Our hypothesis was that human blood cells irradiated *in vivo* in the mouse would show a similar response to that of *ex vivo* irradiated human blood cells.

## Results

### Radiation response of blood cells in human v mouse microarray studies

From our previous studies using microarrays, we have accumulated a substantial amount of data from independent studies on radiation effects on gene expression in blood cells in both human and mouse models. In the course of these studies, we have noticed that some genes appeared to respond to radiation in an opposite direction in mouse and human studies (e.g. up regulated in humans and down regulated in mice). We were, therefore, interested in comparing gene expression between these two species to determine the extent of these differences, as well as better defining the similarities between the two species in order to improve the translatability of radiation signatures between mice and humans. We started by using the datasets summarized in Table 1, which allowed us to compare similar doses in human samples both *in vivo* (3.75 Gy)^18^ and *ex vivo* (8 Gy)^3^; and mouse *in vivo* samples at 4 Gy^23^ and 8 Gy^31^. After identifying the genes that were differentially regulated and comparing these side-by-side between the two species (Figure 1, annotated tables shown in Supplementary table S1), we selected genes to validate the radiation response patterns, and to test the suitability of the hematopoietically humanized mouse as a model for irradiation of human cells *in vivo*. We chose to test this model because it provides an *in vivo* environment for human cells that allows flexibility of experimental design and extension of studies of radiation response to a few days or weeks, which is not possible in *ex vivo* studies. In the current study we focused on the 24 hour response to radiation.

**Table 1.** Summary of microarray studies from which data was compiled for comparisons.

| **Data Set** | **Number of genes**<br>**(p<0.001, FDR\* <5%)** | **GEO accession**<br>**No.** | **Reference** |
| --- | --- | --- | --- |
| **Human 3.75 Gy** | 1588 | GSE20162 | Paul et al <sup>18</sup> |
| **Mouse 4 Gy** | 3577 | GSE85323 | Broustas et al <sup>23</sup> |
| **Human 8 Gy** | 272 | GSE8917 | Paul et al <sup>3</sup> |
| **Mouse 8 Gy** | 780 | GSE99176 | Rudqvist et al <sup>31</sup> |

\* Benjamini-Hochberg false discovery rate

**Figure 1.**
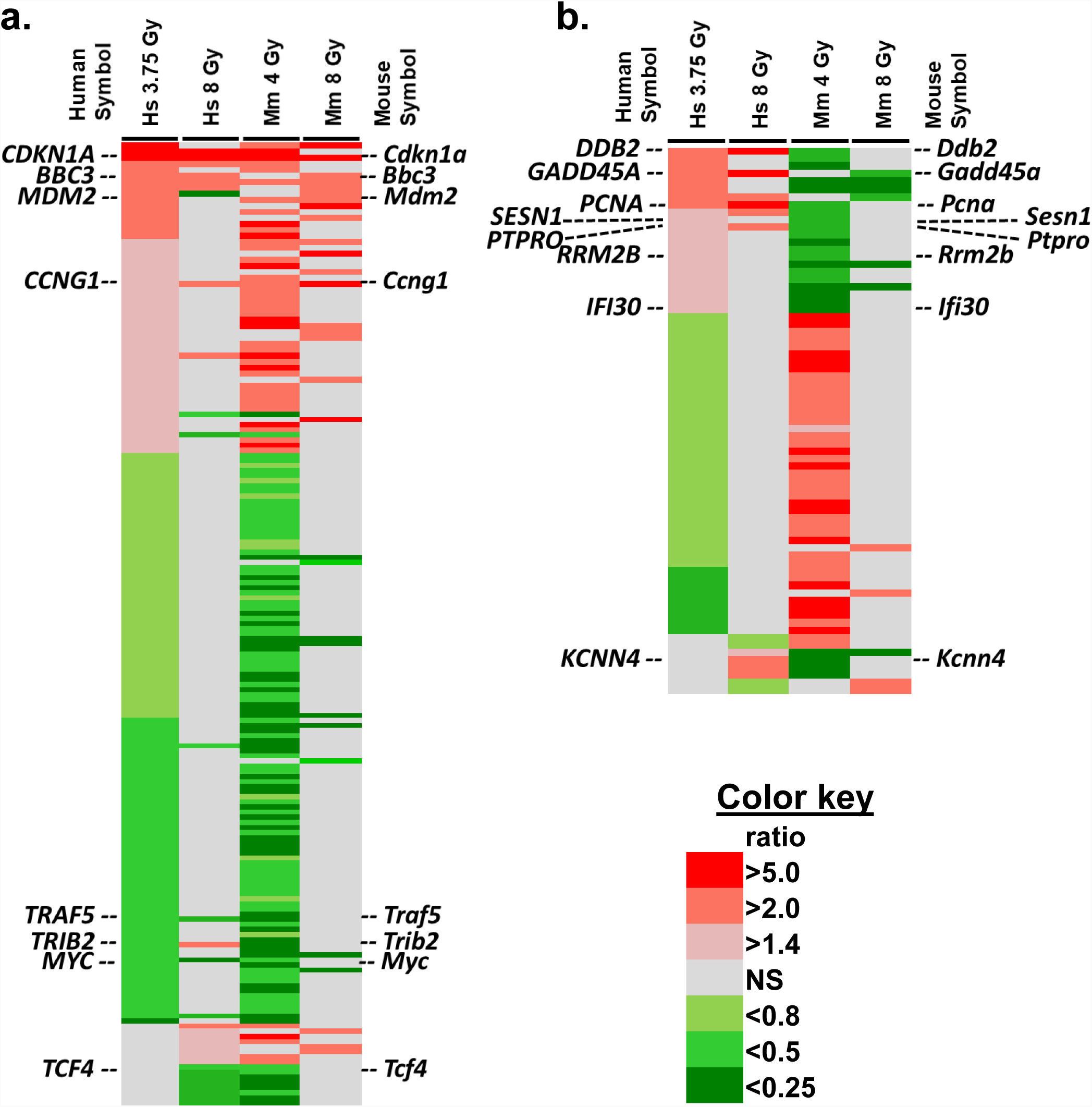
Heat map of gene expression changes from human and mouse microarray studies. a. **Similar response patterns in human and mouse.** The heat map shows homologous genes with similar responses in human and mouse blood after irradiation (doses as shown in Table 1). b. **Opposite responses in human and mouse.** The heat map shows homologous genes with opposite responses in human and mouse blood after radiation (doses as shown in Table 1). Genes selected for qRT-PCR studies in humanized NSG radiation studies are indicated. The key indicates the relative ratios between irradiated and sham-irradiated controls, shades of red for up-regulated genes and shades of green for down-regulated genes. Grey indicates no significant differential expression. A more detailed fully annotated view of these heat maps is shown in Supplementary table S1.

We used the Ingenuity Pathway Analysis software to identify potential upstream regulatory proteins implicated in the discordant responses between species. We based our analyses on the genes that were significantly differentially expressed in opposite directions after 4 Gy in mice ^23^ and 3.75 Gy in humans^18^. Based on this analysis, we determined that TP73 and TP53, related members of the TP53 family of transcriptional regulatory proteins known to play a major role in radiation response, were highly implicated in the discordant radiation response in the two species (Figure 2 and also see Supplementary fig S3 for networks with overlay of fold changes). Additionally MYC, ZBTB16, STAT4 and KLF4 transcriptional regulatory proteins showed this opposite prediction of activity, potentially inactivated in human and activated in mouse blood cells, based on these selected genes.

**Figure 2.**
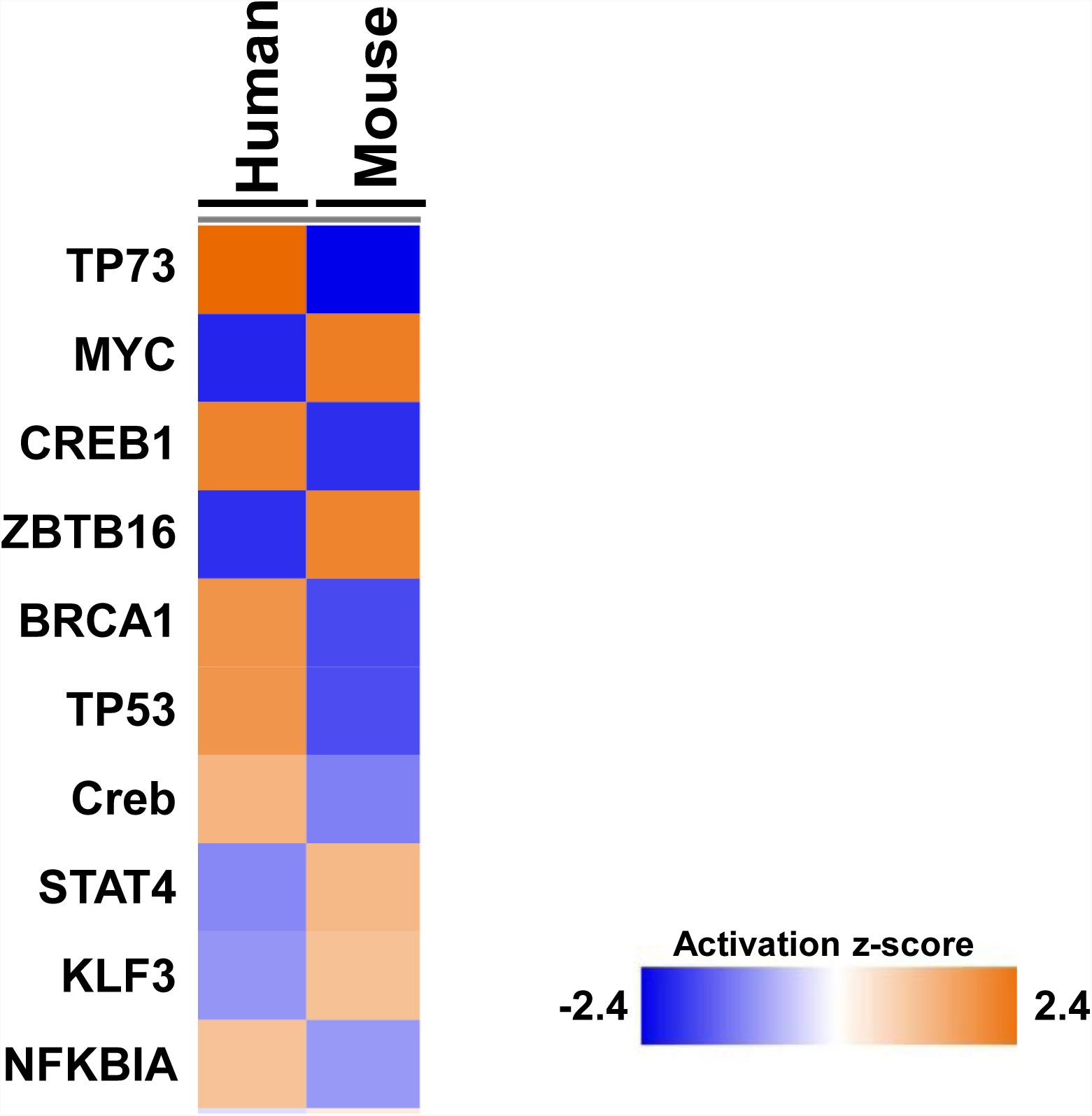
Heat map of predicted transcriptional regulators upstream of opposite microarray gene expression changes. All regulatory proteins shown here were predicted to be involved in the regulation of the set of genes showing opposite activities in humans and mice at ∼4 Gy dose of radiation. The deeper the color (blue for inactivation and orange for activation as determined by the z-score) the stronger the predicted activation/inactivation status of the regulatory protein, as shown in the key.

### Cellular response to radiation

We whole-body irradiated humanized NSG mice at ∼8 months of age (∼5 months after engraftment), as well as age-matched C57BL/6 and non-engrafted NSG mice with sham, 2 Gy and 4 Gy x-rays, and collected blood 24 hours later. C57BL/6 mouse blood gene expression comparisons provided a positive control for repeatability of the previous results from microarrays. In parallel with the mouse samples, we exposed human donor blood to the same doses of x-rays *ex vivo* and collected the blood 24 hours later as a positive control for the human responses.

A portion of the blood samples was used for immunophenotyping, and the results are summarized in Figures 3 and 4. Although there was a trend of decrease in cell density in the blood of humanized mice for human CD45+ (reduced by 53% after 2 Gy and 64% after 4 Gy) and a similar decrease in human CD3+ T cells (reduced by 76% after 2 Gy and 80% after 4 Gy) and human CD19+ B cells (reduced by 80% after 2 Gy and 84% after 4 Gy), Figure 3A; only the decreases in T and B cells from human blood donors irradiated *ex vivo* were statistically significant. *Ex vivo* irradiated human blood samples showed changes that were of lower magnitude, but less variable than the *in vivo* results; CD3+ T cells (reduced 16% after 2 Gy and 29% after 4 Gy, p-value <0.05) and CD19+ B cells (reduced 25% after 2 Gy and 39% after 4 Gy, p-value <0.05), Figure 3B.

**Figure 3.**
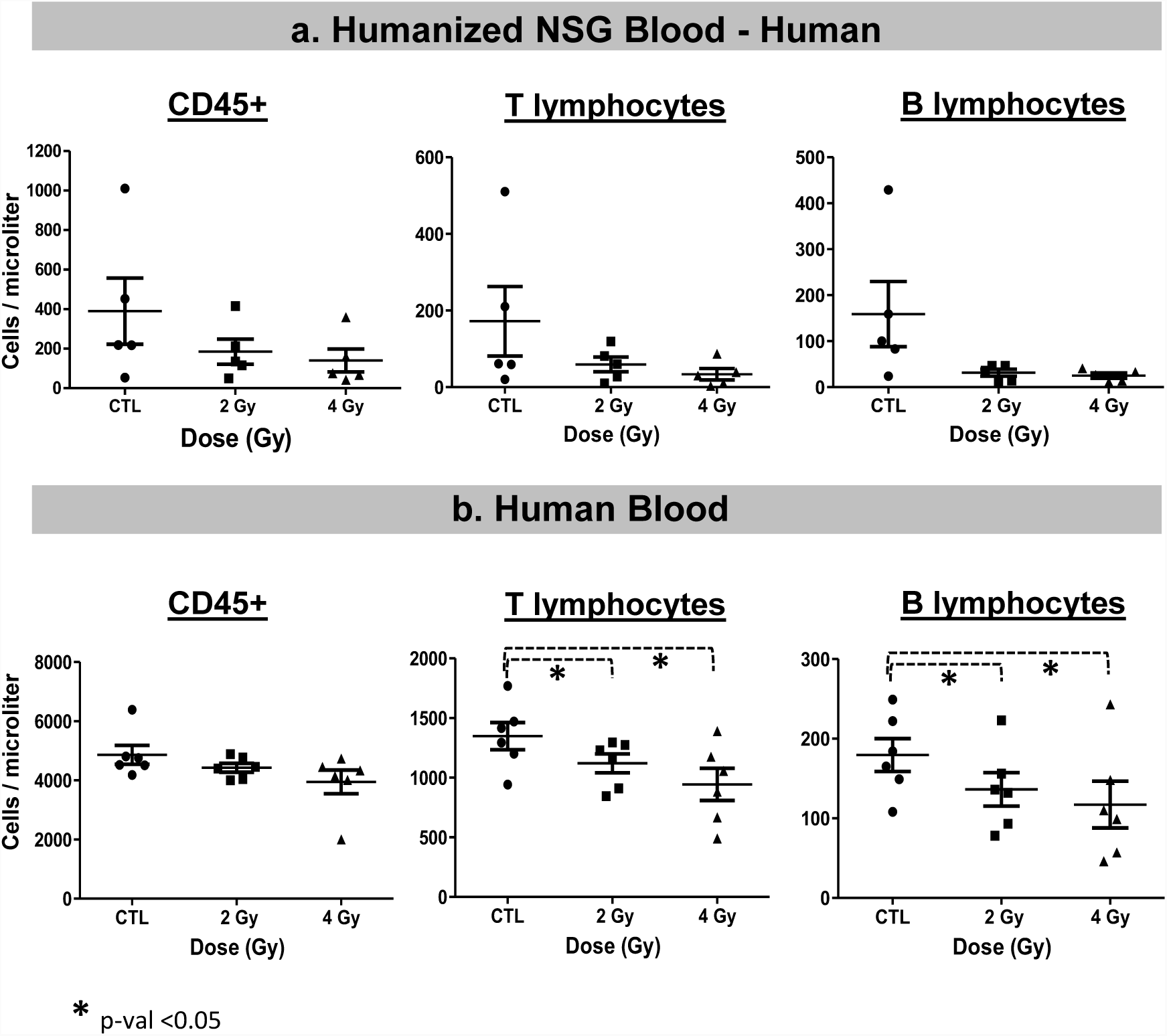
Immunophenotyping of human cells in a. Humanized NSG mice, and b. Human blood 24 hours after *ex vivo* irradiation with 2 and 4 Gy doses. We used a two-tailed t-test (paired for human blood and unpaired for humanized NSG blood) to evaluate statistical significance and * indicates a p-value <0.05.

**Figure 4.**
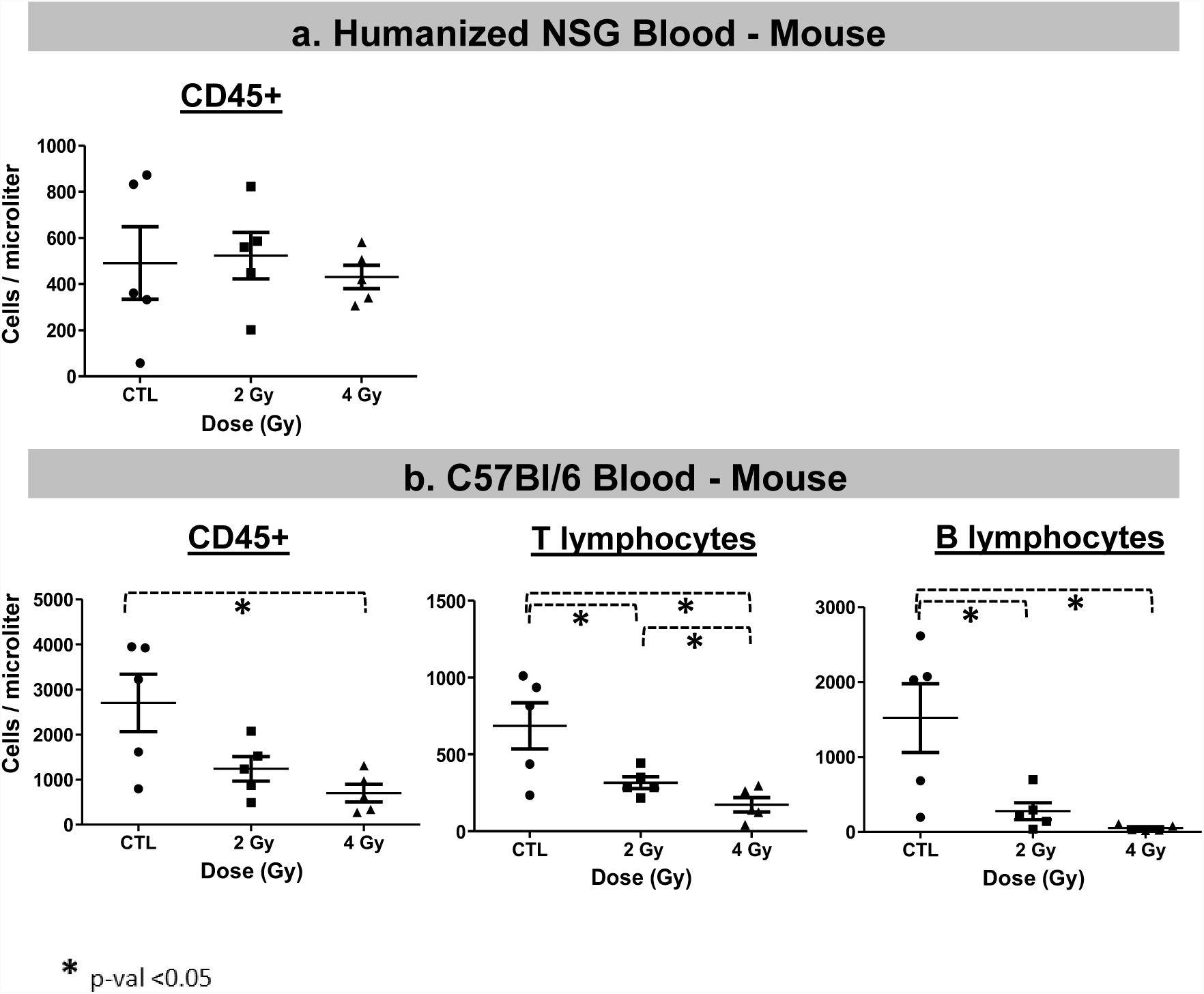
Immunophenotyping of mouse cells in a. Humanized NSG mice, and b. C57BL/6 mice 24 hours after irradiation with 2 and 4 Gy doses. We used an unpaired two-tailed t-test to evaluate statistical significance and * indicates a p-value <0.05.

Mouse blood cells in humanized NSG were present at lower density than in the C57BL/6, and numbers were not affected by the 2 Gy dose. They were only slightly, but not significantly, different after 4 Gy (average cell density reduced by 12% after 4 Gy, p-value >0.05), Figure 4A. In C57BL/6 positive control mouse blood cells, immunophenotyping results showed significant changes after irradiation, Figure 4B. Overall CD45+ lymphocytes were decreased after 2 Gy (reduced by 54%) and 4 Gy (reduced by 74%, p-value <0.05). CD3+ T cells were significantly decreased (reduced by 54% after 2 Gy and 75% after 4 Gy, p-value <0.05); and CD19+ B cells, the most radiosensitive cells, showed a similar response (reduced 82% after 2 Gy and 96% after 4 Gy, p-value <0.05).

### Differential gene expression response to radiation by quantitative real-time PCR

To verify the results of the comparisons between species and their radiation response, we tested the expression of select genes by quantitative PCR analyses in humans and mice and treated humanized NSG mice with the same doses and time point to test the radiation response of human blood cells from this model system. We selected genes for testing by quantitative PCR, based on their response in human v mouse blood cells. We chose the genes by directionality of response, either similar in humans and mice or opposite. We reverse transcribed globin-depleted total RNA from the four experimental groups (Humanized NSG, Human blood *ex vivo*, C57BL/6, and NSG background), after 2 Gy and 4 Gy doses of radiation and performed PCR reactions using Low Density Arrays and determined relative gene expression using sham-irradiated samples as calibrators. These results are shown in Figures 5 and 6, for genes that have similar responses in both species and those that have opposing responses to radiation, respectively.

**Figure 5.**
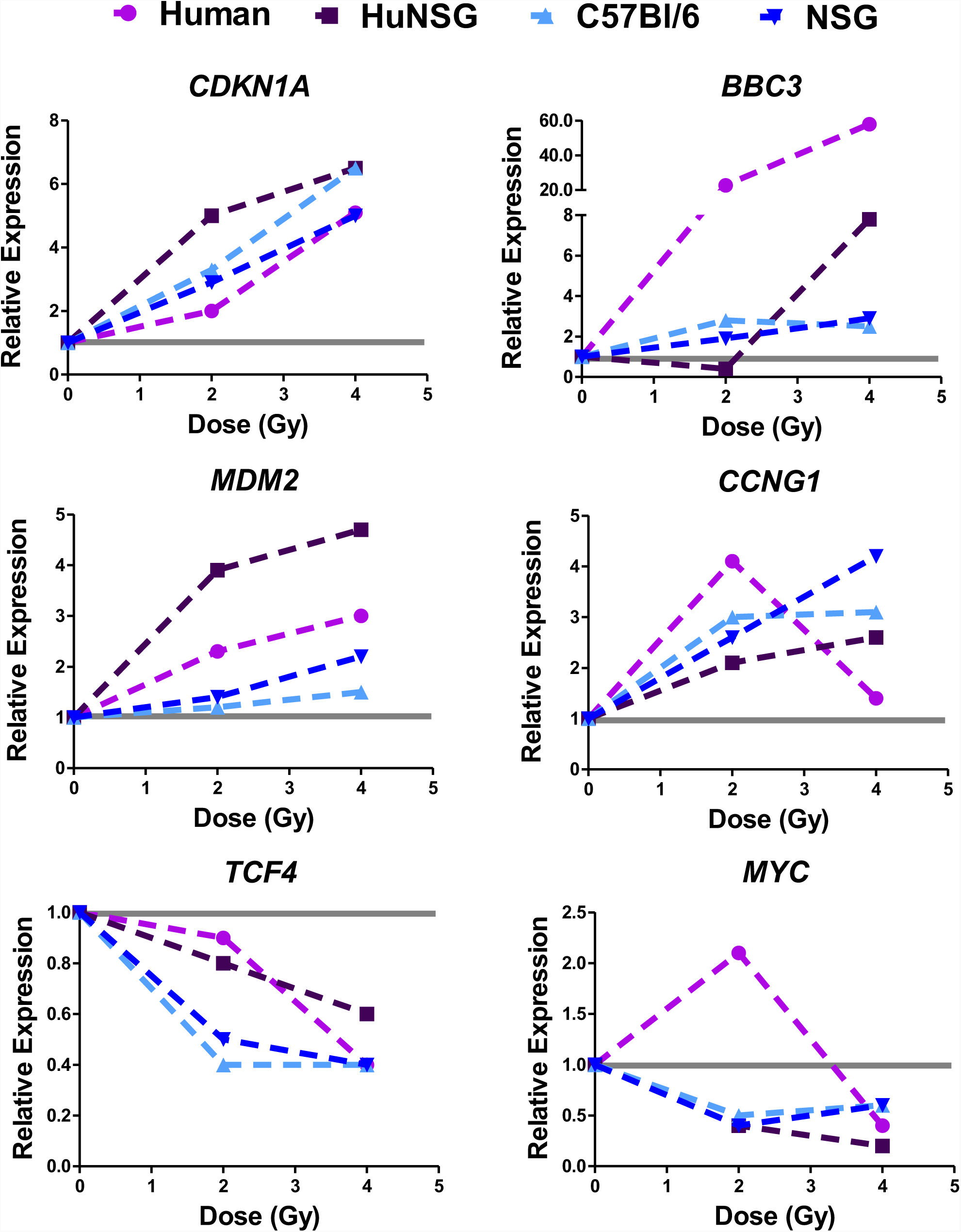
Relative gene expression changes after radiation in human and mouse cells 24 hours after irradiation (similar response patterns). Genes were selected from microarray analyses for showing similar responses to radiation in both human and mouse blood. Values are mean fold change of n>5 biological replicates, for four experimental groups Human (Human blood irradiated *ex vivo*, pink line and symbols), HuNSG (Humanized NSG mice whole body irradiation, human genes, purple line and symbols), C57BL/6 (mice after whole body irradiation, light blue line and symbols) and NSG (age-matched NSG non-engrafted mice after whole body irradiation, dark blue line and symbols). The grey line indicates the calibrator sample, which is the matched sham-irradiated sample for each experimental group.

**Figure 6.**
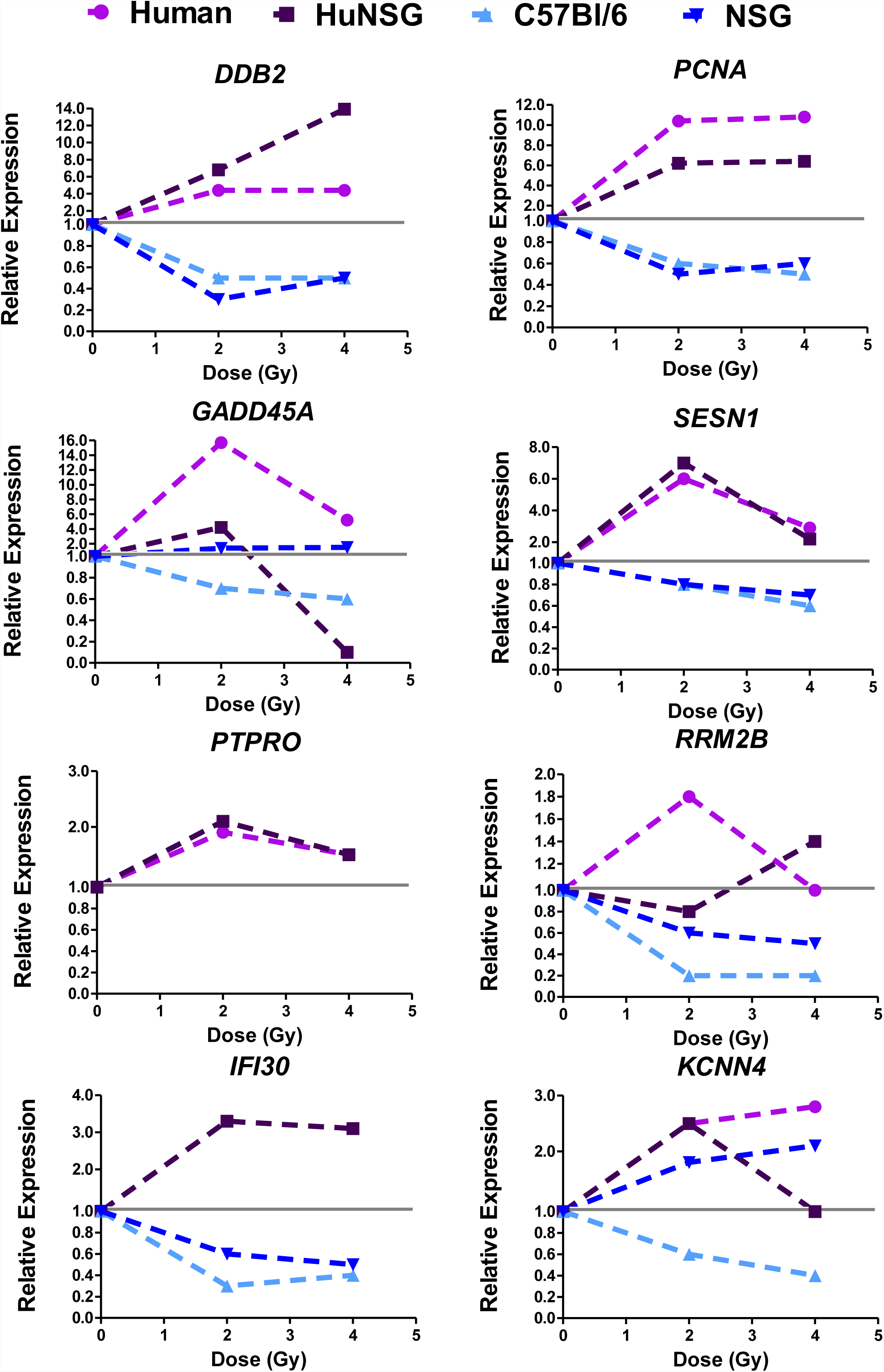
Relative gene expression changes after radiation in human and mouse cells 24 hours after irradiation (opposite response patterns). Genes were selected from microarray analyses for showing opposite responses (at least in one condition) to radiation in human and mouse blood. Values are mean fold change of n>5 biological replicates, for four experimental groups Human (Human blood irradiated *ex vivo*, pink line and symbols), HuNSG (Humanized NSG mice whole body irradiation, human genes, purple line and symbols), C57BL/6 (mice after whole body irradiation, light blue line and symbols) and NSG (age-matched NSG non-engrafted mice after whole body irradiation, dark blue line and symbols). The grey line indicates the calibrator sample, which is the matched sham-irradiated sample for each experimental group. For the *PTPRO* gene, the relative expression of human genes is shown as it agreed with the response as measured by microarrays, up-regulated after radiation, however, the levels of this mRNA were undetectable by qPCR in the mouse samples, C57BL/6 and NSG, therefore are not shown. Most of the NSG transcripts (dark blue) for the selected genes followed the expected C57BL/6 response (light blue) except for the *KCNN4* gene which was up-regulated, as in the human response.

Genes that were selected because of similarity of response to radiation were *CDKN1A, BBC3, MDM2, CCNG1, TCF4* and *MYC*, representing both up and down regulated genes. Genes selected for opposing responses to radiation were *DDB2, PCNA, GADD45A, SESN1, PTPRO, RRM2B, IFI30* and *KCNN4*. Most of these genes have been previously identified in multiple studies as radiation response genes^3,32-50^ however, genes that have similar responses across species have been the primary focus of such investigations. For most similar responding genes in Figure 5, the relative magnitude of change in mRNA levels appears to be closer within the same species, with more difference in the magnitude of response between human and mouse, despite similar trends. Opposite responding genes, showed a directional change in at least one condition tested, so although not all genes showed differential gene expression after radiation at all doses or models tested, the most consistent pattern was that of a divergence between species in the response to radiation.

## Discussion

The relevance of the laboratory mouse as a model organism for investigations of human health and disease is a much discussed issue within the scientific community. The mouse model (*Mus musculus*) has been extensively used and proven extremely helpful in understanding the mechanisms underlying immunological development and processes related to disease and cancer through genetic mutants and transgenic models. However, there is evidence that there are inter-species differences that confound translatability of results from mouse to human, especially as a model for pre-clinical testing of drugs^51,52^. There has been much debate on the advantage of this animal model and in 2013 Seok et al^7^ published a comprehensive study comparing gene expression in blood cells after acute inflammatory stress in which cross-species similarities were lower than expected. The authors proposed that as a result of different biological stimuli during the evolution of mouse and human immune systems, as well as the large differences in life span between the two species, the sensitivity of the mouse and human immune systems differs greatly, which may lead to greater heterogeneity in the murine response. Therefore they also suggested that information about any disease condition in humans, such as the degree of transcriptomic changes, should be tested on a case-by-case basis and may need to be adjusted for in the mouse model used to mimic the human condition. There is also evidence that at the molecular level of genes and transcripts there exist many inter-species differences that have the potential to affect biological responses in different ways^53-57^.

In the field of radiation biology, mice have been used extensively for mortality studies, cytogenetics, and in the last few decades for development of exposure biomarkers or biodosimetric signatures for use after a radiological accident or other large-scale event^2,21^. The search for gene expression biomarkers of radiation exposure has one overall goal, to identify potential gene targets that can be used in a device to rapidly distinguish between unexposed and exposed individuals relative to a threshold dose, or to predict radiation exposure levels. Studies from our group and others have shown that human and mouse homologs of radiation response genes can respond similarly and work very well to predict dose in these two species^2,3,21^. However, we have observed some surprising differences in gene expression response between the species and we designed the present study to address the extent of this divergence.

To start, we compared microarray data to identify patterns and response trends among genes in humans and mouse. We based our meta-analysis on results from four independent studies in which we had irradiated and corresponding sham-irradiated sample groups after exposure of blood to similar doses in the two species. We matched significantly differentially expressed gene homologs at two doses, ∼4 Gy and 8 Gy, and identified 179 unique genes with a similar pattern of response in mouse and humans for at least one dose, and 73 unique genes with an opposite pattern of response for at least one dose (Supplementary table S1). Intriguingly, many of the genes showing an opposite response to radiation were known radiation response genes, and most were up-regulated in humans and down-regulated in mouse. We confirmed the two expression patterns, similar or opposite direction of response, in human and mouse blood cells using qRT-PCR. Finally, we tested the responses in a humanized mouse model to investigate if the human direction of response would be retained in the engrafted human cells.

The initial hematopoietically humanized mouse was able to support maturation of human blood cells for 4-6 months after engraftment, but then would succumb to graft-versus-host-disease (GVHD)^58^. The NSG mouse model used in our study (NSG background: NOD.Cg-Prkdc^scid^ Il2rγ^tm^1^Wjl^/SzJ), has the advantage of improved engraftment levels with human blood CD34+ stem cells^29,30,59,60^. It also does not manifest rejection of the human cells until after >6 months^5,61^, providing a longer window of opportunity for performing experiments. Previous studies using this humanized model to study the effects of radiation have focused on long term effects on the bone-marrow niche, where irradiated human cells showed higher levels of γH2AX foci at 12 weeks after irradiation^62^. The same group also identified down-regulation of B lymphocyte related mRNA after irradiation, which supported the reduction of mature human B lymphocytes and a myeloid-biased population^63^. To our knowledge, our study is the first to address the application of the hematopoietically humanized mouse model to study the early effects of radiation and compare them side-by-side with human and non-engrafted mouse blood cells, to evaluate its usefulness as modified model organism for human stress response.

In our study, we irradiated humanized NSG animals with 0 Gy, 2 Gy and 4 Gy x-rays, and 24 hours after irradiation performed gene expression analysis and cell counts. Hematology results using immunophenotyping and flow cytometry of human cells in the humanized mice did not show dramatic differences in sensitivity between human T and B cells after radiation in the peripheral blood (Figure 3A). Although all HuNSG in this study were engrafted with stem cells from one donor^58^, the broad range of engraftment efficiency could contribute to heterogeneity between sub-populations after irradiation. Also, the doses in this study were 2 Gy and 4 Gy, which are relevant doses for biomarker discovery for triage after radiological events, but may be extremely toxic in NSG mice, which are radiation sensitive^5,29,30^. In human blood cells after *ex vivo* irradiations, as expected B cells were more sensitive to radiation (Figure 3B, 16% depletion after 2 Gy in T cells and 25% depletion of B cells after the same dose)^64^. Mouse lymphocytes from C57BL/6 were also sensitive to radiation, with B cells being more sensitive than T cells (Figure 4B, 54% depletion after 2 Gy in T cells and 82% depletion of B cells after the same dose)^64^. The hematological results presented here indicate that, despite documented differences in the immune systems of mice and humans (reviewed by Mestas et al^52^), the humanized NSG mouse model is sensitive to radiation, and the *in vivo* microenvironment it provides for engrafted human cells promotes rapid removal of damaged lymphocytes and may present a more relevant context than *ex vivo* exposures ^65^.

The patterns of radiation response of genes selected from the meta-analysis of human v mouse blood microarray data were all confirmed in our qRT-PCR experiments (Figures 5 and 6). Moreover, the gene expression response to radiation in human cells from HuNSG mice followed closely with human TBI or *ex vivo* blood responses, indicating that the cellular milieu of human cells in the mouse blood does not affect the radiation gene response, via inter-cellular signaling from mouse to human.

We also used IPA upstream analysis to suggest transcriptional regulatory proteins that may contribute to the inter-species differences observed here (Figure 2). TP53, TP73, MYC, CREB and STAT transcription factors were predicted to be potential upstream regulators of the set of genes showing the opposite pattern of response to radiation, based on the gene expression from microarrays. Due to its central role in radiation response, we focused on TP53. All of the genes in Figure 5 are known target genes of the TP53 transcriptional regulation network that respond similarly in mouse and human. This emphasizes the fact that TP53 is clearly activated in response to radiation exposure in both species. However, TP53 regulated genes such as *DDB2, PCNA, GADD45A* and *SESN1* were down-regulated by irradiation in the mouse, in contrast to their up-regulation in humans (Figure 6). IPA identified 23 genes from the meta-analysis of radiation response that showed this opposite response to radiation and that had a direct molecular interaction with TP53 (Supplementary fig S3). The overlay of relative expression values for these 23 genes (15 up regulated and 8 down regulated in human, and *vice versa* in mouse), suggests that human and mouse p53 may have different effects on the expression of a subset of their target genes, perhaps on the basis of different post-translational modifications or different avidity or availability of regulatory co-factors.

The TP53 transcription factor is a central player in many biological processes, including stress response and disease. It can undergo modifications at the DNA, RNA and protein level in order to perform a variety of biological functions. Degradation of TP53 is a well-known feedback mechanism to control its activity^66^. Structurally, the TP53 protein has many modifications that can dictate its function and cellular fate ^67^ and recent investigations show differences in the dynamics of TP53 activation across different mammalian species^68,69^. In fact, a comparison of TP53 response elements (REs or TP53 binding sites) using a reporter assay across 14 species revealed that the transactivation of target genes in mice (related to functions such as cell cycle, apoptosis, DNA metabolism and angiogenesis) was only moderate in comparison to the response of human REs^70^. Another study in which global TP53 transactivation analyses were performed to identify target genes in embryonic stem cells (ESC), it was revealed that the TP53-transcriptional program may have undergone evolutionary divergence leading to developmental functions in human ESC and DNA damage functions in the mouse^71,72^. A comparison of single cell dynamics of TP53 oscillations (negative feedback via MDM2, which is also transcriptionally induced by TP53, leading to oscillations in TP53 levels) in four mammalian species showed that in humans, non-human primates and canines, TP53 oscillations are slower than in mouse and that this is due to differences in the balance between TP53 degradation and MDM2 transcription rates^68^. These regulatory circuits and others can be altered in response to radiation and may contribute to the differences in our results presented here. Although we do observe conservation of patterns in certain genes (Figure 5) due to the broad redundancy in stress response transcription factors species-specific differences in transcriptional activity may affect only a small set of downstream target genes (Figure 6).

The results of this study emphasize the need to consider the impact of signaling differences between species in translational studies. Also, effects of other species-specific mechanisms such alternative splicing of regulators, localization of regulatory proteins, redundancy of transcriptional regulation, etc. cannot be ruled out as contributing to the opposite response observed here. However, since these species differences are intra-cellular, and appear not to be communicated between cells or from the microenvironment, by using hematopoietically humanized mice, we were able to largely recapitulate the human radiation response in an *in vivo* context, adding a powerful tool for future mechanistic studies and also for studies of long-term radiation response.

Our study began as a result of observations of differences in the results of global transcriptomics analyses of radiation response in human and mouse blood cells. A more extensive analysis of the literature comparing more species, doses and times may reveal additional genes that show differences between species. Future studies to address these genes and their impact on biomarker discovery will expand our knowledge of the differences between humans and mouse and most importantly will aid in the application of mouse biological responses to human health. These results also strongly support the use of humanized mice as a useful model organism for translational studies of radiation response and biomarker discovery.

## Methods

### Comparison of human and mouse radiation gene expression

In order to compare the gene expression response in mouse and human, we used data from our previously published gene expression studies of radiation responses in mice and humans at 24 hours post exposure. Differentially expressed gene lists from the blood of human total body irradiation (TBI) patients (3.75 Gy delivered in 3 × 1.25 Gy fractions) or *ex vivo* exposed peripheral blood (8 Gy) were compared with those from mice exposed to 4 or 8 Gy x-rays, all at 24 hours post exposure (see Table 1). Human *in vivo* exposure was represented by the samples from patients undergoing TBI, with ratios representing the response to 3.75 Gy delivered in 3 fractions over 24 hours^18^ which was compared with a 4 Gy exposure in mice^23^. In order to extend the comparison to a higher dose, we included data from human donor blood exposed to 8 Gy *ex vivo*^3^ and from mice also exposed to 8 Gy^31^. We built a relational database in FileMaker® Pro and populated it from the human-murine homolog file downloaded from Mouse Genome Informatics^73^ on 27^th^Feb2018. We imported expression ratio values from the four data sets by matching on the human or mouse gene name, and then extracted the data from the homologs responding significantly in both mouse and human (Supplementary table S1). These gene lists and the gene expression values in human and mouse blood after irradiation were used for network analysis using Ingenuity Pathway Analysis and for identification of genes for further analysis in this study.

### Network analysis

We uploaded the lists of human and mouse differentially expressed genes to Ingenuity Pathway Analysis® Software (IPA from Ingenuity® http://www.ingenuity.com) and performed prediction analysis for upstream regulators, which not only identifies potential upstream regulators and ranks them by statistical significance but also provides information about the direction of activation of that regulatory protein based on the downstream gene targets from the gene list. The program provides a z-score that uses the number of target genes and the type of relationship between the regulator and target genes (either activation or inhibition) from the published literature.

### Engraftment of NSG mice with human blood stem cells

All animal husbandry and experimental procedures were conducted in accordance with applicable federal and state guidelines and approved by the Animal Care and Use Committees of Columbia University (Assurance Number: A3007-01). C57BL/6NCrl mice were purchased from Charles River Laboratories (Wilmington, MA). Immuno-deficient NSG (NOD.Cg-Prkdc^scid^ Il2rg^tm^1^Wjl^/SzJ) mice (The Jackson Laboratory; Bar Harbor, ME), aged 5 to 7 weeks, were engrafted with commercially available human cord blood CD34+ cells (Cincinnati Children’s Hospital Medical Center, Cincinnati, OH, USA). For engraftment, the NSG mice were irradiated with 2 Gy of γ-rays using a Gammacell-40 ^137^Cs irradiator (AECL, Ottawa, Ontario, Canada), followed by injection of 200,000 human CD34+ cells into the mouse tail vein within 24 hour after the irradiation. Twelve weeks later, ∼50 microliters of blood was obtained from each mouse (all male) by tail artery bleeding, and engraftment was quantified by standard flow cytometry methods, using antibodies specific to human blood cell surface antigens for CD45 (Biolegend, catalog# 103115), CD3 (Biolegend, catalog# 100312), CD20 (Biolegend, catalog# 115508), CD11b (Biolegend, catalog# 101210) and mouse blood cell surface antigen CD45 (Biolegend, catalog# 103115). Engraftment of human blood cells in the humanized NSG mice used in this study ranged from 20% to 74% human lymphocytes.

### Irradiations, collection of blood and RNA isolation

All mice and human blood samples were irradiated in groups of 5 using an X-RAD Biological Irradiator (Precision X-ray, North Branford, CT) at a dose rate of 1.03 Gy/min at a machine setting of 320 KeV/12.5 mA. Doses were sham, 2 Gy and 4 Gy. 24 hours after irradiation mice were rapidly euthanized by CO_2_ asphyxia and blood was collected using a 21-gauge syringe via cardiac puncture. The volume was recorded, an aliquot was taken for immunophenotyping analyses as described above, and the remaining blood was mixed into PAXgene® solution (Becton Dickinson, NJ) in a 1:5 ratio and inverted >10 times to mix. Immunophenotyping analysis of the blood human and mouse cell subpopulations was done by flow cytometry using the antibodies mentioned in the previous subsection. After storage of blood in PAXgene® tubes 4°C overnight, RNA was isolated from whole blood using the PAXgene® Blood RNA kit (Qiagen, Valencia, CA) according to the manufacturer’s protocol. Human blood samples were collected with informed consent from healthy donors following the guidelines and regulations of the Institutional Review Board protocol of Columbia University Medical Center (IRB-AAAF2671). Peripheral blood samples were collected in Sodium Citrate vacutainer tubes (Becton Dickinson, NJ, catalog# 366415) and irradiated. Immediately after irradiation the human peripheral blood samples were mixed with culture media (RPMI + 10% heat inactivated FBS with 1% penicillin/streptomycin) in 50 mL Tube Spin Bioreactor® tubes (TTP, Switzerland) in a tissue culture incubator at 37°C and 5% CO_2_ for 24 hours, before RNA was isolated using the PerfectPure RNA isolation kit (5 Prime, Gaithersburg, MA). The mouse and human RNA samples were globin-depleted using the species-specific Ambion® GLOBINclear kit (Ambion, Thermofisher) and the RNA was quantified using a NanoDrop One spectrophotometer (Thermofisher).

### Quantitative real time PCR analysis of human and mouse genes

We designed Low Density Arrays (LDA) using Taqman® assays from Life Technologies/ Thermofisher as listed in Supplementary table S2. The selection of the Taqman® primer-probe assays was based on *in silico* tests of cross-reactivity, based on sequence matching of the primers between species on the ThermoFisher gene search website (https://www.thermofisher.com). We experimentally validated all human assays for non-cross-reactivity with mouse cDNA and *vice versa* testing the mouse assays against human cDNA. These tests verified that none of the PCRs gave Ct values <33 which we consider to be the minimum sensitivity of detection for Taqman® assays.

We prepared complimentary DNA (cDNA) from total mRNA using the High-Capacity® cDNA Kit (Life Technologies, Foster City, CA). Quantitative real-time RT-PCR (qRT-PCR) was performed for the selected genes using Taqman® assays (Life Technologies) on LDA cards. For the humanized NSG mouse group, 1000 ng of cDNA was used as input for human assays, while 250 ng cDNA was used as input for the mouse assays compensating for the fraction of human: mouse material in the cDNA. The increased input of cDNA from humanized NSG mice for human assay reactions was based on preliminary testing of the sensitivity of the assays in the humanized mouse samples, which consist of mixed human and mouse cDNA. We tested two different amounts of cDNA input: 250 ng and 1000 ng and found that increased cDNA input was required to detect human housekeeping gene transcripts at a similar Ct value to those from using 250 ng of the same cDNA pool for mouse housekeeping gene transcripts. For single species reactions, human or mouse, 250 ng cDNA input was used. Quantitative real time PCR reactions were performed with the ABI 7900 Real Time PCR System using Universal PCR Master Mix (Thermofisher), with initial activation at 50°C for 120 seconds and 95°C for 10 minutes, followed by 40 cycles of 97°C for 30 seconds and 59.7°C for 60 seconds. Relative fold-induction was calculated by the -ΔΔCT method^74^ using SDS version 2.4 (Thermofisher). Data were normalized to the geometric mean of 5 human and mouse housekeeping genes (*ACTB, GAPDH, UBC, YWHAZ*, and *HPRT1*), which were found to be stably expressed across all samples in human and mouse mRNA pools using geNorm^75^.

## Declarations

### Ethics and approval and consent to participate

For animal work: All animal husbandry and experimental procedures were conducted in accordance with applicable federal and state guidelines and approved by the Animal Care and Use Committees of Columbia University (Assurance Number: A3007-01). For human work: Human blood samples were collected with informed consent from healthy donors following the guidelines and regulations of the Institutional Review Board protocol of Columbia University Medical Center (IRB-AAAF2671)

### Availability of data and material

Microarray datasets used for meta-analyses in this study are publically available in the NCBI Gene Expression Omnibus database (https://www.ncbi.nlm.nih.gov/geo/) under accession numbers GSE20162, GSE85323, GSE8917 and GSE99176.

## Acknowledgements

We would like to acknowledge Ms. Mashkura Chowdhury and Ms. Aesis Luna for assistance with mouse handling and irradiations.

## Authors’ contributions

SAG and SAA designed the study, SAG performed the experiments, analyzed the data, and drafted the manuscript. SAA helped with preparation of the manuscript. MP and LBS engrafted the humanized mice and contributed to the manuscript. All authors have read and approved the final manuscript.

## Additional Information

## Supplementary Information

accompanies this paper.

## List of Supplementary files

**Supplementary table S1** Gene expression comparison between human and mouse

**Supplementary table S2** Low Density Array design

**Supplementary figure S3** Ingenuity networks of TP53 in humans and mouse

